# Two engineered AAV capsid variants for efficient transduction of human cortical neurons directly converted from iPSC

**DOI:** 10.1101/2021.09.06.459070

**Authors:** Sandra Fischer, Jonas Weinmann, Frank Gillardon

**Affiliations:** CNS Diseases Research, Boehringer Ingelheim Pharma GmbH & Co. KG, 88397 Biberach an der Riss, Germany

## Abstract

Recombinant adeno-associated virus (AAV) is the most widely used vector for gene therapy in clinical trials. To increase transduction efficiency and specificity, novel engineered AAV variants with modified capsid sequences are evaluated in human cell cultures and non-human primates. In the present study, we tested two novel AAV capsid variants, AAV2-NNPTPSR and AAV9-NVVRSSS, in human cortical neurons, which were directly converted from human induced pluripotent stem cells and cocultured with rat primary astrocytes. AAV2-NNPTPSR variant efficiently transduced both induced human cortical glutamatergic neurons and induced human cortical GABAergic interneurons. By contrast, AAV9-NVVRSSS variant transduced both induced human cortical neurons and cocultured rat primary astrocytes. High viral titers (1×10E5 viral genomes per cell) caused a significant decrease in viability of induced human cortical neurons. Low viral titers (1×10E4 viral genomes per cell) lead to a significant increase in the neuronal activity marker c-Fos in transduced human neurons following treatment with a potassium channel blocker, which may indicate functional alterations induced by viral transduction and/or transgene expression.

## 1. Introduction

Recombinant adeno-associated virus (AAV) vectors have been widely used in clinical trials for gene therapy of neurological disorders (reviewed in (Iqubal et al., 2020; Piguet et al., 2021)). In most clinical studies AAV vectors based on naturally occurring AAV serotypes were applied, which exhibit different transduction efficiency and cell tropism due to variations in the AAV capsid sequence. To increase transduction efficiency and specificity, novel AAV variants were engineered by modification of the AAV capsid sequence (reviewed in (Grimm and Büning, 2017; Kotterman and Schaffer, 2014)). Barcoded libraries of different AAV capsid variants from independent screening campaigns were subsequently screened in mice to identify superior vectors for human gene therapy (Weinmann et al., 2020; Westhaus et al., 2020). However, mouse models lack human-specific cellular signatures, which may be important for AAV transduction. Screening of AAV capsid variants in different human primary cell types already pointed to peptide motifs resulting in enhanced transduction rates of specific cell types (Börner et al., 2020; Westhaus et al., 2020). Human neurons differentiated from induced human pluripotent stem cells (iPSC) are increasingly used as a test system for AAV vectors designated for gene therapy of neurological disorders (Depla et al., 2020; Duong et al., 2019). Neuronal differentiation of human iPSC by small molecules however, is a lengthy procedure and leads to a heterogeneous mixture of various neuron types and non-neuronal cells (Strano et al., 2020; Yao et al., 2017). By contrast, lentiviral overexpression of neuralizing transcription factors rapidly converts human iPSC into a homogenous population of mature cortical neurons (Nehme et al., 2018; Zhang et al., 2013). In the present study, we generated induced human cortical glutamatergic neurons and induced human cortical GABAergic interneurons, respectively, to test the transduction efficiency of two engineered AAV vectors that have shown improved transduction rates of other human cell types.

## 2. Materials and methods

### 2.1. Culture of human iPSC

Human iPSC were cultured as described by us in detail previously (Fischer et al., 2020). In short, iPSC were seeded in mTeSR medium on culture plates coated with Matrigel basement membrane matrix. Cells were split using 0.02% EDTA and were replated in mTeSR medium supplemented with 10□µM ROCK inhibitor Y27632. Human iPSC had a normal karyotype and were negatively tested for mycoplasma (Fischer et al., 2020).

### 2.2. Direct neuronal conversion of iPSC

Human iPSC were directly converted into glutamatergic cortical neurons by lentiviral overexpression of the neuralizing transcription factor Ngn2. Alternatively, iPSC were converted into GABAergic cortical interneurons by lentiviral overexpression of the transcription factors Ascl1 and Dlx2, as originally described by others (Wang et al., 2017; Zhang et al., 2013). For accelerated neural induction, direct neuronal conversion was combined with developmental pattering by pharmacological inhibition of TGF-beta, BMP, and WNT signaling as published by others (Nehme et al., 2018; Qi et al., 2017). Induced neurons were cocultured with rat primary cortical astrocytes on day 8 post induction to promote neuronal maturation and synapse formation (Zhang et al., 2013). Neuronal function/activity was assessed by multi-electrode array recordings and whole-cell patch-clamp recordings (Fig. 1) (Fischer et al., 2020). Synaptic activity-induced c-Fos expression was triggered by administration of the potassium channel blocker 4-aminopyridine at 100 µM for 3h (Pruunsild et al., 2017).

**Fig. 1.**
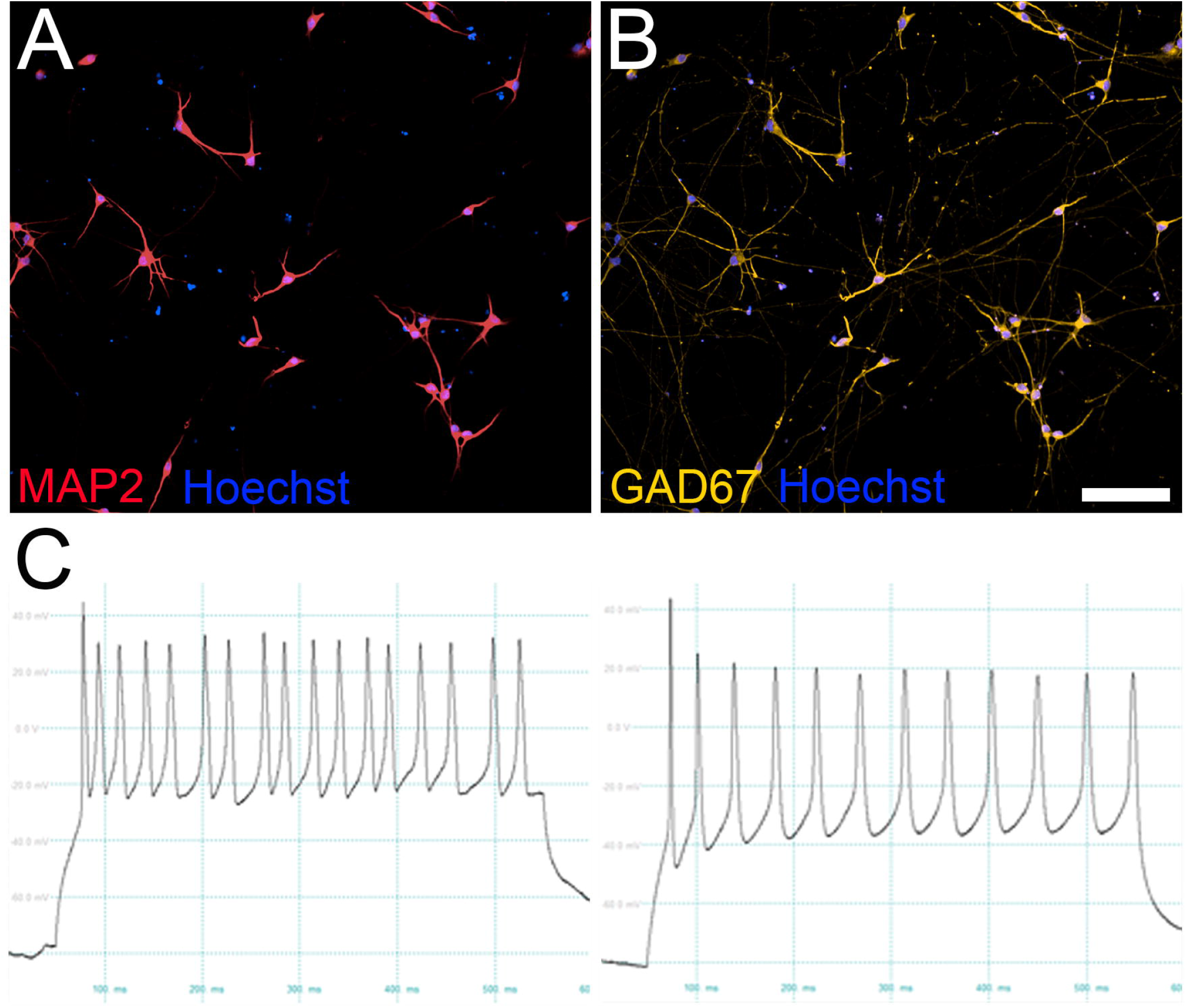
(A, B) Representative immunofluorescence microscopy images of human cortical interneurons directly converted from human iPSC by Ascl1 and Dlx2 overexpression. Cell cultures were immunostained for MAP2, a pan-neuronal marker, and for GAD67, a marker for GABAergic interneurons. Most nuclear Hoechst 33324-labeled cells are immunopositive for both MAP2 and GAD67. Scale bar represents 100 µm. (C) Representative whole-cell patch-clamp recordings from two iPSC-derived GABAergic interneurons show sustained action potential firing following current injection.

### 2.3. AAV production and transduction of cell cultures

Cloning, production, purification, and titration of AAV vectors were performed as described in detail previously (Börner et al., 2020; Strobel et al., 2019; Völkner et al., 2021; Weinmann et al., 2020). Expression of the enhanced green fluorescent protein (GFP) reporter was driven by the ubiquitously active cytomegalovirus promotor. We selected two second generation AAV capsid variants for testing. Capsid variant AAV2-NNPTPSR has been shown by others to efficiently transduce various cell types in human retinal explants and human retinal organoids (Pavlou et al., 2021; Völkner et al., 2021). Capsid variant AAV9-NVVRSSS was selected based on a high transduction efficiency in AAV library screens in human cells focusing on the NXXRXXX peptide motif (Börner et al., 2020) (Weinmann et al., unpublished data). By contrast, wildtype AAV2 and AAV9 serotypes poorly transduce human iPSC-derived neurons (Depla et al., 2020; Duong et al., 2019) (Weinmann et al., unpublished data). Cocultures of induced human neurons and primary rat astrocytes in 96-well plates were transduced with AAV vectors encoding an enhanced GFP reporter driven by the ubiquitously active cytomegalovirus promotor at day 21 of maturation. AAV vectors were added at 1×10E4 viral genomes per cell or 1×10E5 viral genomes per cell. These titers are in the lower range of previous studies, where commonly used AAV serotypes were tested in human forebrain neurons differentiated from iPSC by small molecules (Depla et al., 2020; Duong et al., 2019). Cell culture medium was exchanged three days after AAV administration. Sham-treated cultures were used as controls.

### 2.4. Immunofluorescence staining and digital image analysis

Immunostaining and image analysis were performed as described by us in detail elsewhere (Fischer et al., 2020; Kizner et al., 2020). In short, cells were fixed in 4% paraformaldehyde. Thereafter, cells were permeabilized in 0.1% Triton X-100, unspecific protein binding was blocked in 5% normal goat serum, and primary antibodies were added over night at 4°C. The following primary antibodies were used for immunostaining: anti-MAP2 (EnCor Biotechnology, Gainesville, USA), anti-vGLUT1 (Synaptic Systems, Goettingen, Germany), anti-GAD67 (Merck, Darmstadt, Germany), anti-GFP (Thermo Fisher Scientific, Waltham, USA), and anti-c-Fos (Sigma-Aldrich, St. Louis, USA) (Fischer et al., 2020). After three washing steps, Alexa-conjugated secondary antibodies were added for 2 hours at room temperature. Hoechst 33342 fluorescent dye was used to stain nuclei. Fluorescent signals were detected using the Opera Phenix High-Content Screening System, and digital images were analyzed using Columbus software. Cells showing non-condensed/non-fragmented, Hoechst 33324-fluorescent nuclei were counted as viable cells (Naujock et al., 2020).

### 2.5. Statistical analysis

For AAV transduction and microscopic analysis, cells were randomly assigned to the cell culture plates. For subsequent data acquisition, investigators were blinded regarding the group category. Graph Pad Prism version 8 (GraphPad Software, San Diego, USA) was used for all statistical analyses. Data are from two independent experiments. All graphs show mean ± standard deviation. Statistical testing was performed using Student’s two-tailed, unpaired t-test. The significance level was set to 5% per hypothesis.

## 3. Results

For phenotypic characterization, human cortical neurons directly converted from iPSC by overexpression of Ngn2 were double immunostained for MAP2, a pan-neuronal marker, and for vGLUT1, a marker for glutamatergic neurons. Human cortical neurons directly converted from iPSC by overexpression of Ascl1 and Dlx2 were double immunolabeled for MAP2 and for GAD67, a marker for GABAergic interneurons. Consistent with published data (Wang et al., 2017; Zhang et al., 2013), more than 90% of the MAP2 stained neurons were vGLUT1-positive, glutamatergic neurons or GAD67-positive, GABAergic interneurons as assessed by digital image analysis (Fig. 1) (Fischer et al., 2020).

Induced human cortical neurons were cocultured with rat primary astrocytes to promote neuronal maturation. Cell cultures from the same batch were transduced with second generation AAV capsid variants AAV2-NNPTPSR or AAV9-NVVRSSS at two different titers (1×10E4 viral genomes per cell or 1×10E5 viral genomes per cell). Transduction efficiency was initially monitored using fluorescence microscopy of living cultures. As shown in Figure 2 A-D, preferential enhanced GFP reporter gene expression in human cortical glutamatergic neurons is visible four days after transduction with AAV2-NNPTPSR capsid variant. By contrast, several GFP-fluorescent cells showing astrocytic/non-neuronal morphology are detected in cultures transduced with AAV9-NVVRSSS capsid variant.

**Fig. 2.**
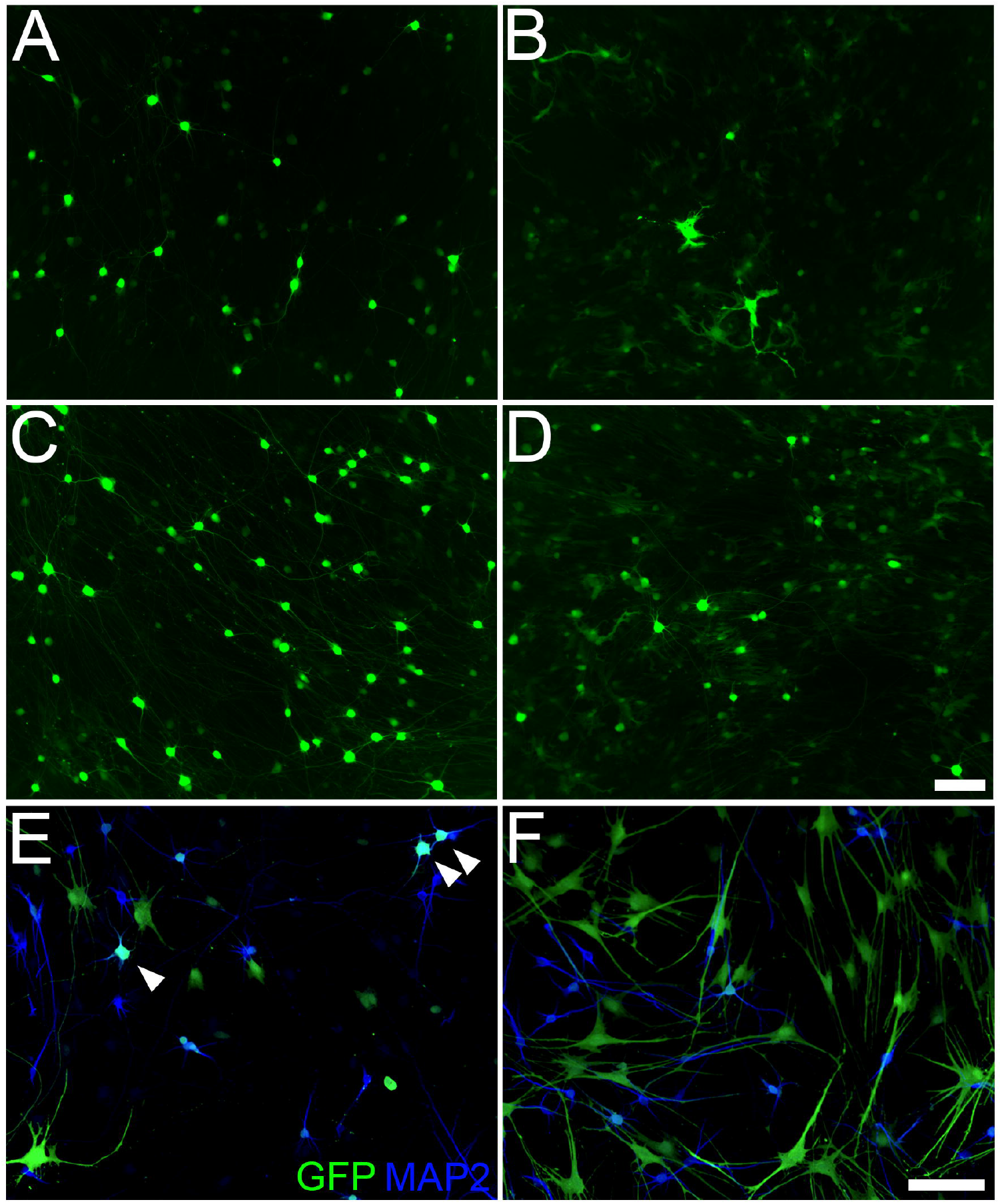
(A-D) Representative fluorescence microscopy images showing AAV-mediated enhanced GFP expression in human cortical glutamatergic neurons directly converted from iPSC by lentiviral overexpression of Ngn2. Induced human neurons were cocultured with rat primary astrocytes. Cell cultures were transduced with AAV capsid variants at 1×10E4 viral genomes per cell (A, B) and at 1×10E5 viral genomes per cell (C, D), respectively. (A, C) Four days post transduction with AAV2-NNPTPSR capsid variant, live imaging revealed preferential enhanced GFP expression in cells showing neuronal morphology. (B, D) By contrast, several GFP-fluorescent cells showing astrocytic/non-neuronal morphology were visible in cultures transduced with AAV9-NVVRSSS capsid variant. (E, F) Representative immunofluorescence images showing AAV-mediated enhanced GFP expression in human cortical GABAergic interneurons directly converted from iPSC by lentiviral overexpression of Ascl1 and Dlx2. Induced human interneurons were cocultured with rat primary astrocytes. Cell cultures were transduced with 1×10E4 viral genomes per cell and were immunostained 8 days post AAV transduction. (E) In cell cultures transduced with AAV2 capsid variant, enhanced GFP expression was observed both in several MAP2-positive interneurons (arrowheads) and in several MAP2-negative, astrocytes/non-neuronal cells. (F) By contrast, exclusively GFP-positive, MAP2-negative transduced astrocytes/non-neuronal cells were visible in cell cultures transduced with AAV9 capsid variant. Scale bars represent 100 µm.

In human cortical GABAergic interneuron cultures transduced with AAV2 capsid variant, GFP immunofluorescence is visible both in human interneurons and in astrocytes/non-neuronal cells, whereas exclusively GFP-immunopositive, transduced astrocytes/non-neuronal cells are detectable in cell cultures transduced with AAV9 capsid variant (Fig. 2 E, F). By microscopic inspection of living cultures, the number of GFP-fluorescent, transduced cells increased up to day 8 after AAV transduction. Therefore, we concentrated on transduction of induced human cortical glutamatergic neurons and on day 8 post transduction in subsequent experiments.

Immunostaining of transduced cells followed by digital image analysis of 96-well plates were used to quantify transduction efficiency, cellular viability, and neuronal function at single cell resolution. As shown in Figure 3, double immunostaining for GFP and the pan-neuronal marker MAP2 confirmed a significant (p < 0.001) and almost exclusive transduction of induced human cortical glutamatergic neurons by AAV2-NNPTPSR capsid variant. By contrast, a significant (p < 0.001) increase both in transduced human neurons and in transduced astrocytes/non-neuronal cells was detected following transduction with AAV9-NVVRSSS capsid variant (Fig. 3 G).

**Fig. 3.**
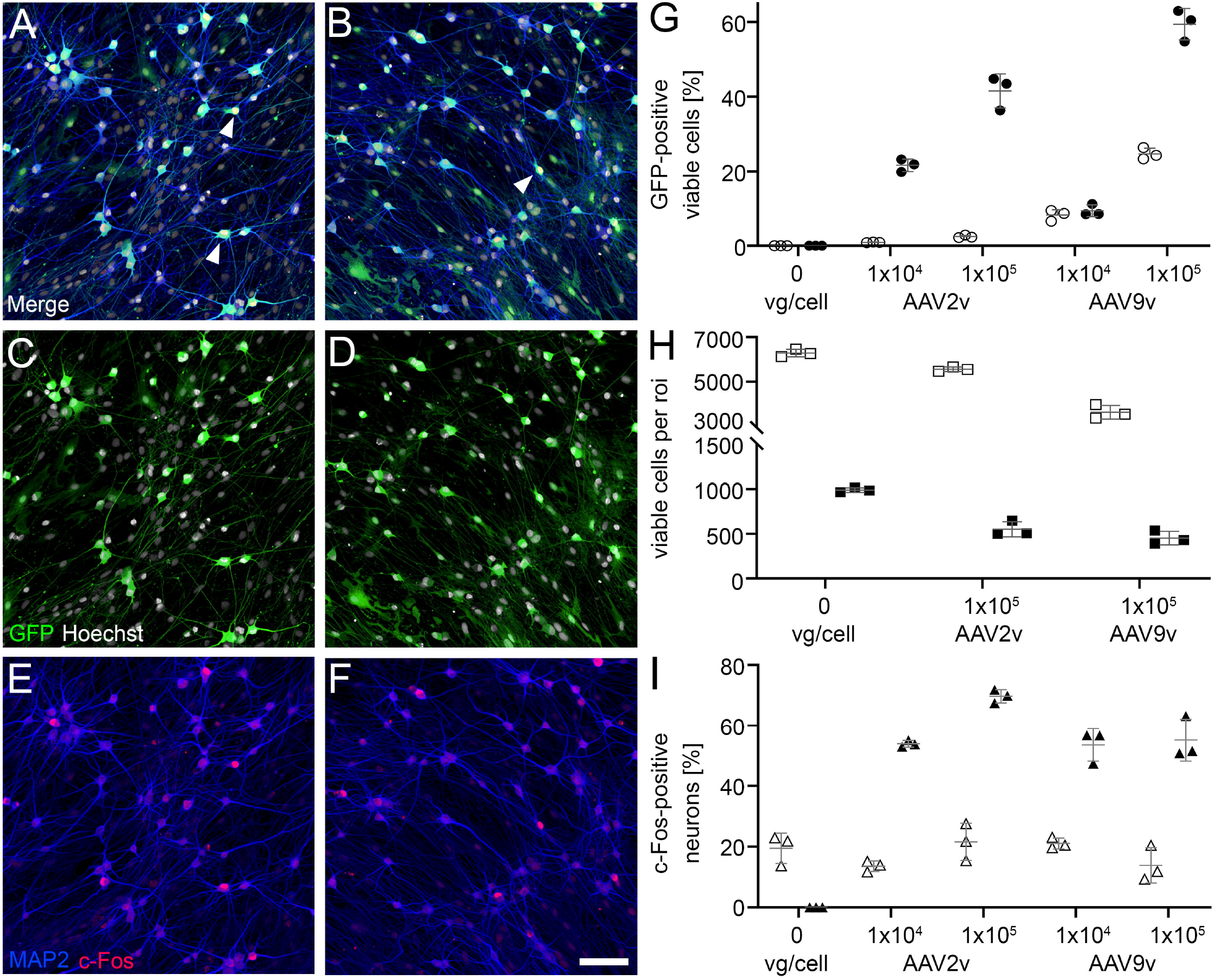
(A-F) Representative microscopy images following immunofluorescent staining of induced human cortical glutamatergic neurons cocultured with rat primary astrocytes for AAV-mediated enhanced GFP expression (green), the pan-neuronal marker MAP2 (blue), and the neuronal activity marker c-Fos (red). Nuclei were labelled with fluorescent Hoechst 33324 dye (white). (A, C, E) Images show cell cultures 8 days after transduction with AAV2-NNPTPSR capsid variant at 1×10E5 viral genomes per cell. (B, D, F) Images show cell cultures 8 days after transduction with AAV9-NVVRSSS capsid variant at 1×10E5 viral genomes per cell. Synaptic activity-induced c-Fos protein expression was triggered by administration of the potassium channel blocker 4-aminopyridine (100 µM, 3h). Scale bar represents 100 µm. (G, H, I) High-content digital image analysis of cell cultures after AAV-transduction and immunofluorescent staining using the Opera Phenix imaging system and Columbus software. Cells showing non-condensed/non-fragmented, Hoechst 33324-fluorescent nuclei (arrowheads in A, B) were counted as viable cells. (G) A significant (p < 0.001) increase in GFP-positive, MAP2-positive, transduced human cortical neurons (filled circles) is detectable 8 days following AAV2 variant (AAV2v) transduction at 1×10E4 viral genomes per cell and at 1×10E5 viral genomes per cell, respectively. By contrast, both GFP-positive, MAP2-positive, transduced human cortical neurons (filled circles) and GFP-positive, MAP2-negative, transduced astrocytes/non-neuronal cells (empty circles) significantly (p < 0.001) increase following transduction with AAV9 variant (AAV9v). (H) After AAV transduction at 1×10E5 viral genomes per cell, a significant decrease in viability of MAP2-positive, human cortical neurons (filled squares, p < 0.001) and of total cells (empty squares, p < 0.01) per region of interest (roi) is detected by digital image analysis. (I) Quantitative analysis of c-Fos-positive, GFP-positive, MAP2-positive transduced human cortical neurons (filled triangles) and c-Fos-positive, GFP-negative, MAP2-positive non-transduced human cortical neurons (empty triangles) in the same well. Nuclear c-Fos protein expression was induced by administration of the potassium channel blocker 4-aminopyridine (100 µM, 3h). A significant (p < 0.01) higher percentage of c-Fos-positive, transduced human cortical neurons compared to c-Fos-positive, non-transduced human neurons is detectable. All figures show mean ± standard deviation. Forty-nine regions of interest per well and three wells per experimental group were analyzed.

Digital image analysis of viable cells showing non-condensed/non-fragmented, Hoechst 33324-fluorescent nuclei revealed a significant decrease in viability of human cortical neurons (p < 0.001) and of total cells (p < 0.01) following AAV transduction at 1×10E5 viral genomes per cell (Fig. 3 H). Transduction at 1×10E4 viral genomes per cell did not affect cell viability.

To assess neuronal function after AAV transduction, the potassium channel blocker 4-aminopyridine was added to the cultures (100 µM, 3h), which significantly increases neuronal spiking activity and nuclear c-Fos expression in iPSC-derived neurons (Pruunsild et al., 2017). Notably, a significant (p < 0.01) higher percentage of c-Fos-positive, GFP-positive, transduced human cortical neurons compared to c-Fos-positive, GFP-negative, non-transduced human neurons is detectable in the same well at both viral titers (Fig. 3 I), which may indicate functional alterations induced by viral transduction and/or transgene expression.

## 4. Discussion

Naturally occurring AAV serotypes have been used in numerous clinical trials for gene therapy of neurological disorders (reviewed in (Iqubal et al., 2020; Piguet et al., 2021)). To increase transduction efficiency of human cells, engineered AAV capsid variants are increasingly tested in human cell cultures and non-human primates. In the present study, we evaluated two novel engineered AAV capsid variants, AAV2-NNPTPSR and AAV9-NVVRSSS. Insertion of the NNPTPSR peptide has recently been shown by others to significantly increase transduction of human retinal neurons and glia both in human retinal explant cultures and in human iPSC-derived retinal organoids (Pavlou et al., 2021; Völkner et al., 2021). By contrast, wildtype AAV2 and AAV9 serotypes poorly transduce human iPSC-derived neurons (Depla et al., 2020; Duong et al., 2019) (Weinmann et al., unpublished data). In our study, AAV2-NNPTPSR variant showed a highly efficient and specific transduction of induced human cortical glutamatergic neurons, whereas AAV9-NVVRSSS variant transduced both induced human cortical glutamatergic neurons and cocultured rat primary astrocytes. Interestingly, we observed an increased transduction of primary rat astrocytes when cocultured with induced human cortical GABAergic interneurons. Since astrocyte gene expression is modulated by neuronal cues (Bayraktar et al., 2020), it may be hypothesized that cues from induced human GABAergic interneurons lead to increased expression of the (unknown) receptors targeted by these AAV variants in cocultured rat astrocytes.

Importantly, we detected a significant decrease in viability of human cortical neurons following AAV transduction at 1×10E5 viral genomes per cell and a significant increase in the neuronal activity marker c-Fos in transduced human neurons already at 1×10E4 viral genomes per cell. Neuronal cell death was not observed in human retinal organoids 10 days after transduction with AAV2-NNPTPSR variant at 5×10E10 viral genomes per organoid (Völkner et al., 2021). By contrast, degeneration of dorsal root ganglion neurons was detected in non-human primates following AAV9 injection into the cerebrospinal fluid, which may be caused by high transgene expression (Hordeaux et al., 2020).

To our knowledge, neuronal function/excitability has not been analyzed in human neuron cultures following AAV transduction in previous studies. The threefold higher percentage of c-Fos positive, transduced human neurons compared to c-Fos positive, non-transduced human neurons in our study sounds a note of caution. Further studies are required to elucidate, whether increased expression of a marker for neuronal activity in AAV transduced, cultured human neurons is indicative of dysfunctional neuronal networks following AAV administration into the brain.

## Acknowledgements

The authors thank Prof. Moritz Rossner for providing lentiviral vectors, and Jessica Lindner for providing patch-clamp data.

